# DeepHBV: A deep learning model to predict hepatitis B virus (HBV) integration sites

**DOI:** 10.1101/2021.01.08.425855

**Authors:** Canbiao Wu, Xiaofang Guo, Mengyuan Li, Xiayu Fu, Zeliang Hou, Manman Zhai, Jingxian Shen, Xiaofan Qiu, Zifeng Cui, Hongxian Xie, Pengmin Qin, Xuchu Weng, Zheng Hu, Jiuxing Liang

**Affiliations:** Key Laboratory of Brain, Cognition and Education Sciences, Ministry of Education, China; Institute for Brain Research and Rehabilitation, South China Normal University, Guangzhou, China.; Department of Medical Oncology of the Eastern Hospital, the First Affiliated Hospital, Sun Yat-sen University, Guangzhou, Guangdong, China; Department of Gynecological Oncology, the First Affiliated Hospital, Sun Yat-sen University, Guangzhou, Guangdong, China; Department of Thoracic Surgery, the First Affiliated Hospital, Sun Yat-sen University, Guangzhou, Guangdong, China; School of Psychology, South China Normal University, Guangzhou, Guangdong, China; Generulor Company Bio-X Lab, Guangzhou, Guangdong, China; Department of Obstetrics and Gynecology, Tongji Hospital, Tongji Medical College, Huazhong University of Science and Technology, Wuhan, Hubei, China

## Abstract

Hepatitis B virus (HBV) is one of the main causes for viral hepatitis and liver cancer. Previous studies showed HBV can integrate into host genome and further promote malignant transformation. In this study, we developed an attention-based deep learning model DeepHBV to predict HBV integration sites by learning local genomic features automatically. We trained and tested DeepHBV using the HBV integration sites data from dsVIS database. Initially, DeepHBV showed AUROC of 0.6363 and AUPR of 0.5471 on the dataset. Adding repeat peaks and TCGA Pan Cancer peaks can significantly improve the model performance, with an AUROC of 0.8378 and 0.9430 and an AUPR of 0.7535 and 0.9310, respectively. On independent validation dataset of HBV integration sites from VISDB, DeepHBV with HBV integration sequences plus TCGA Pan Cancer (AUROC of 0.7603 and AUPR of 0.6189) performed better than HBV integration sequences plus repeat peaks (AUROC of 0.6657 and AUPR of 0.5737). Next, we found the transcriptional factor binding sites (TFBS) were significantly enriched near genomic positions that were paid attention to by convolution neural network. The binding sites of AR-halfsite, Arnt, Atf1, bHLHE40, bHLHE41, BMAL1, CLOCK, c-Myc, COUP-TFII, E2A, EBF1, Erra and Foxo3 were highlighted by DeepHBV attention mechanism in both dsVIS dataset and VISDB dataset, revealing the HBV integration preference. In summary, DeepHBV is a robust and explainable deep learning model not only for the prediction of HBV integration sites but also for further mechanism study of HBV induced cancer.

**Author summary:** Hepatitis B virus (HBV) is one of the main causes for viral hepatitis and liver cancer. Previous studies showed HBV can integrate into host genome and further promote malignant transformation. In this study, we developed an attention-based deep learning model DeepHBV to predict HBV integration sites by learning local genomic features automatically. The performance of DeepHBV model significantly improves after adding genomic features, with an AUROC of 0.9430 and an AUPR of 0.9310. Furthermore, we enriched the transcriptional factor binding sites of proteins by convolution neural network. In summary, DeepHBV is a robust and explainable deep learning model not only for the prediction of HBV integration sites but also for the further study of HBV integration mechanism.

## Introduction

HBV is the main cause of viral hepatitis and liver cancer (hepatocellular carcinoma: HCC) [1]. It is a small DNA virus that can integrate into the host genome via an RNA intermediate [1]. First, HBV attaches and enters into hepatocytes, then transports its nucleocapsid which contains a relaxed circular DNA (rcDNA) to the host nucleus. In host nucleus, rcDNA is converted into covalently closed circular DNA (cccDNA) which produces messenger RNAs (mRNA) and pregenomic RNA (pgRNA) by transcription. Via reverse transcription in host nucleus, pgRNA produces new rcDNA and double-stranded linear DNA (dslDNA), which tend to integrate into the host cell genome [2].

Previous study showed HBV integration breakpoints distributed randomly across the whole genome with a handful of hotspots [3]. For instance, HBV was reported to recurrently integrate into the telomerase reverse transcriptase (TERT) and Myeloid/lymphoid or mixed-lineage leukemia 4 (MLL4, also known as KMT2B) genes. The insertional events were also accompanied by the altered expression of the integrated gene [2,3,5], indicating important biological impacts on the local genome. Further analysis revealed that the association between HBV integration and genomic instability existed in these insertional events [4]. Moreover, significant enrichment of HBV integration was found near the following genomic features in tumours compared to non-tumour tissue: repetitive regions, fragile sites, CpG islands and telomeres [2]. However, the pattern and the mechanism of HBV integration still remained to be explored. Many of the HBV integration sites distributed throughout the human genome and seem completely random [4,6,7]. Whether the features and patterns of these “random” viral integration events could be learned and extracted remained an open question, and once solved, will greatly improve the understanding towards HBV integration induced carcinogenesis.

Deep learning has an excellent performance in computational biology research, such as medical image identification [8], discovering motifs in protein sequences [9]. The convolutional neural network (CNN) is the most important part in deep learning, which enables a computer to learn and program itself from training data [10]. Though deep learning performs excellent in a various of fields, the detailed theory of how it makes the decision was hard to explain due to its black box effect. Therefore, an approach named attention mechanism which can highlight the outstanding parts was invented to open the “black box” [11,12].

In this study, we developed, DeepHBV, an attention-based model to predict the HBV integration sites using deep learning. The attention mechanism calculates the attention weight for each position and connect the encoder and the decoder in the meanwhile. It highlights the regions concentrated by DeepHBV and helps figure out the patterns that were paid attention to. DeepHBV can predict HBV integration sites accurately and specifically, and the attention mechanism identified positions with potential important biological meanings.

## Results

### DeepHBV effectively predicts HBV integration sites by adding genomic features

DeepHBV model structure and the scheme of encoding a 2 kb sample into a binary matrix were described in Fig 1. DeepHBV model was tested with our HBV integration sites database (http://dsvis.wuhansoftware.com). HBV integration sequences were prepared according to HBV integration sites as positive/negative samples following the steps in Method. The negative samples should be twice number of positive samples to keep data balance and to improve the confidence level. The positive samples were divided into 2902 and 1264 as positive training dataset and testing dataset. Ccorrespondingly, we extracted 5804 and 2528 negative samples as negative training dataset and testing dataset. DeepHINT, an existing deep learning model for predicting HIV integration sites according to surroundings [15], will also be evaluated using HBV integration sequences for training and testing. Both models were trained by the same HBV integration training dataset and used the same testing dataset for the evaluation. DeepHBV with HBV integration sequences showed an AUROC of 0.6363 and an AUPR of 0.5471 while DeepHINT with HBV integration sequences demonstrated an AUROC of 0.6199 and an AUPR of 0.5152 (Fig 2). The comparison of DeepHBV and DeepHINT was described in **Discussion** part.

**Figure 1.**
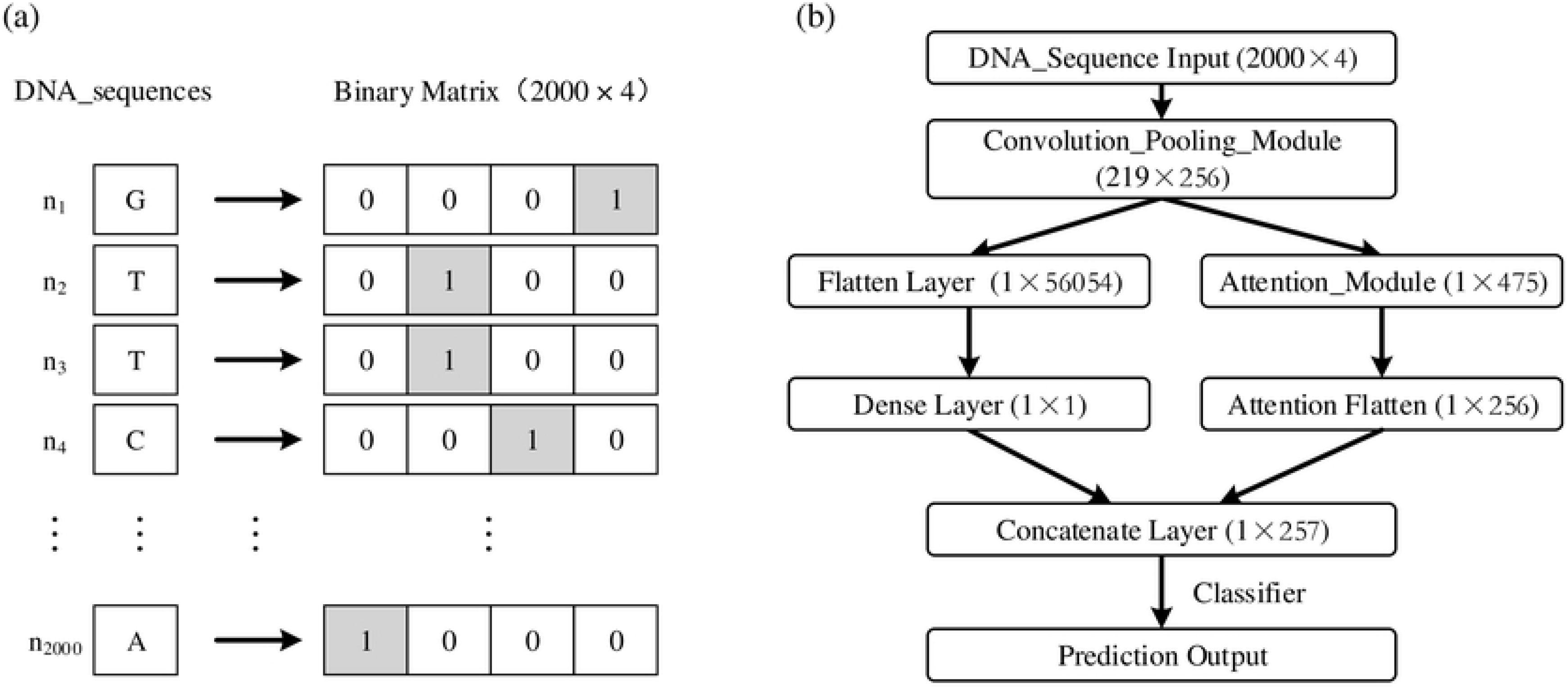
The deep learning framework applied in DeepHBV. (a) Scheme of encoding a 2 kb DNA sequence into a binary matrix using one-hot code; (b) A brief flowchart of DeepHBV structure, the matrix shape was included in brackets, and a detailed flowchart was in S1 Fig.

**Figure 2.**
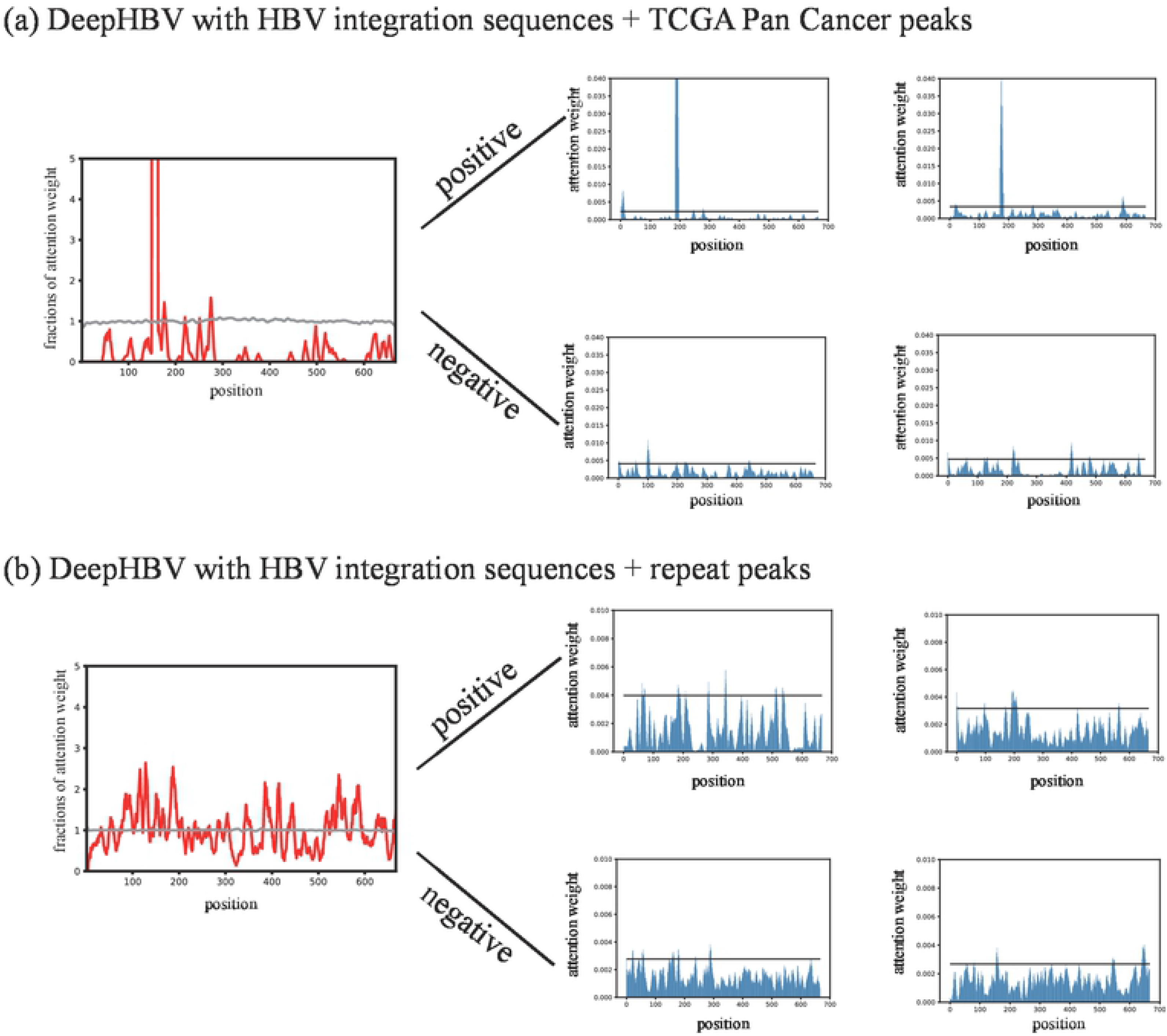
Evaluation of DeepHBV and DeepHINT model prediction performance on the test dataset. (a) receiver-operating characteristic (ROC) curves and (b) precision recall (PR) curves, respectively. “DeepHBV with HBV integration sequences” refers to DeepHBV model with only HBV integration sequences as input; “DeepHINT with HBV integration sequences” refers to DeepHINT model with only HBV integration sequences as input; “DeepHBV with HBV integration sequences + repeat” refers to DeepHBV integration sequences and repeat sequences as input; “DeepHBV with HBV integration sequences” refers to DeepHBV integration sequences and TCGA Pan Cancer sequences as input: “DeepHBV with HBV integration sequences + repeat + (test) VISDB” refers to DeepHBV using HBV integration sequences and repeat sequences for training and using VISDB as independent test dataset; “HBV with HBV integration sequences + TCGA Pan Cancer + (test) VISDB” refers to DeepHBV using HBV integration sequences as TCGA Pan Cancer sequences for training and using VISDB as independent test dataset.

Several previous studies showed that HBV integration has a preference on surrounding genomic features such as repeat, histone markers, CpG islands, etc [2,4]. Thus, we tried to add these genomic features into DeepHBV, by mixing genomic feature samples together with HBV integration sequences as new datasets, then trained and tested the updated DeepHBV models. We downloaded following genomic features from different datasets [16–18] into four subgroups: (1) DNase Clusters, Fragile site, RepeatMasker; (2) CpG islands, GeneHancer; (3) Cons 20 Mammals, TCGA Pan-Cancer; (4) H3K4Me3 ChIP-seq, H3K27ac ChIP-seq (S2 Fig). After obtaining genomic feature data positions (sources are mentioned in S2 Table), we extended the positions to 2000 bp and extracted related sequences on hg38 reference genome. We defined these sequences as positive genmoic feature samples. Then we mixed HBV integration sequences, positive genome feature samples, and randomly picked negative genomic feature samples (see **Method**) together and trained the DeepHBV model. Once a subgroup performed well, we re-test each genomic feature in that subgroup to figure out which specific genomic feature affect the model performance significantly (S2 Fig) (AUROC and AUPR values were recorded in S3 Table). From the ROC and PR curves, we found DeepHBV with HBV integration sites plus the genomic features repeat (AUROC: 0.8378 and AUPR: 0.7535) and TCGA Pan Cancer (AUROC: 0.9430 and AUPR: 0.9310) can significantly improve the HBV integration sites prediction performance against DeepHBV with HBV integration sequences (Fig 2). We also performed the same test on DeepHINT, but did not find a subgroup can substantially improve the model performance (results were recorded in S3 Table). Together, DeepHBV with HBV integration sequences plus repeat or TCGA Pan Cancer can significantly improve the model performance.

### Validation of DeepHBV using independent dataset VISDB

It is necessary of DeepHBV to be applied on general datasets, we tested the pre-trained DeepHBV models (DeepHBV with HBV integration sequences + repeat peaks and DeepHBV with HBV integration sequences + TCGA Pan Cancer peaks) on the HBV integration sites dataset in another viruses integration sites (VIS) database VISDB [19]. We found that in the model trained with HBV integration sequences + repeat sequences showed an AUROC of 0.6657 and an AUPR of 0.5737, while the model trained with HBV integrated sequences + TCGA Pan Cancer showed an AUROC of 0.7603 and an AUPR of 0.6189.

The DeepHBV model with HBV integration sequences + TCGA Pan Cancer performed better compared with DeepHBV model with HBV integration sequences + repeat and was more robust on both testing dataset from dsVIS (AUROC: 0.9430 and AUPR: 0.9310) and independent testing dataset from VISDB (AUROC: 0.7603 and AUPR: 0.6189). Thus, we decided to use this model for future HBV integration sites study.

### Study the preference pattern of HBV integration by conserved sequence elements

DeepHBV can extract features with translation invariance by pooling operation, which enables DeepHBV to recognise certain patterns even the features were slightly translated. The participating of attention mechanism into DeepHBV framework might partly open the deep learning black box by giving an attention weight to each position. Each attention weight represented the computational importance level of that position in DeepHBV judgement. The attention weights in attention layer were extracted after two de-convolution and one de-pooling operation and the output shape is 667×1. Each score represented an attention weight of a 3 bp region. Positions with higher attention weight scores might have more important impact on the pattern recognition of DeepHBV, meaning these positions might be the critical points for identifying HBV integration positive samples. We first averaged the fractions of attention scores in all HBV integration sequences and normalized them to the mean of all positions. Then we visualised the fractions of attention scores and found the figure showed peak-valley-peak patterns only in positive samples (Fig 3). We were interested in the positions with higher attention weights in convolution neural network. And we found that, in the attention weight distribution of DeepHBV with HBV integration sites + TCGA Pan Cancer, a cluster of attention weights much higher than other weights often occurred in the positive samples. While in the model of DeepHBV with HBV integration sites + repeat did not show this pattern (Fig 3).

**Figure 3.**
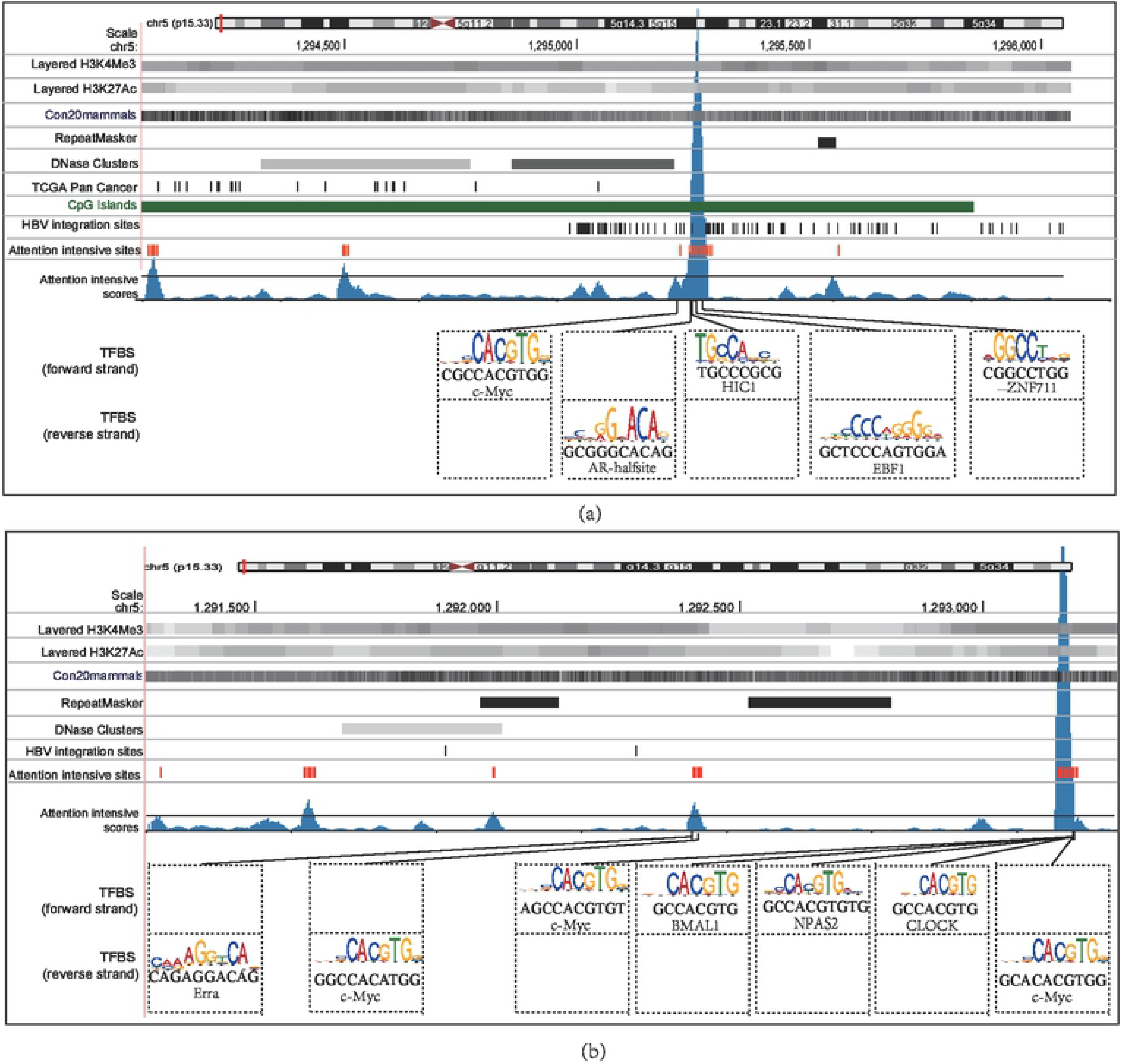
The attention weight distribution of analysed by DeepHBV with HBV integration sequences + genomic features. (a) DeepHBV with HBV integration sequences + TCGA Pan Cancer peaks; (b) DeepHBV with HBV integration sequences + repeat peaks. The left graph showed the fractions of attention weight, which were averaged among all samples and normalized to the average of all positions, each index represents a 3 bp region due to the multiple convolution and pooling operation. The graphs on the right are representative samples of attention weight distribution of positive samples and negative samples.

To further discover the pattern behind these positions with higher attention weights, we defined the sites with top 5% highest attention weights as attention intensive sites, the regions of 10 bp near them as attention intensive regions. We mapped these attention intensive sites on hg38 reference genome with genomic features (Fig 4), but found that the positional relationship between attention intensive sites and genomic features was not quite clear. The results indicated that there may exist other specific pattern closely related to HBV integration preference, and when analysed carefully, could be recognized by the DeepHBV model.

**Figure 4.**
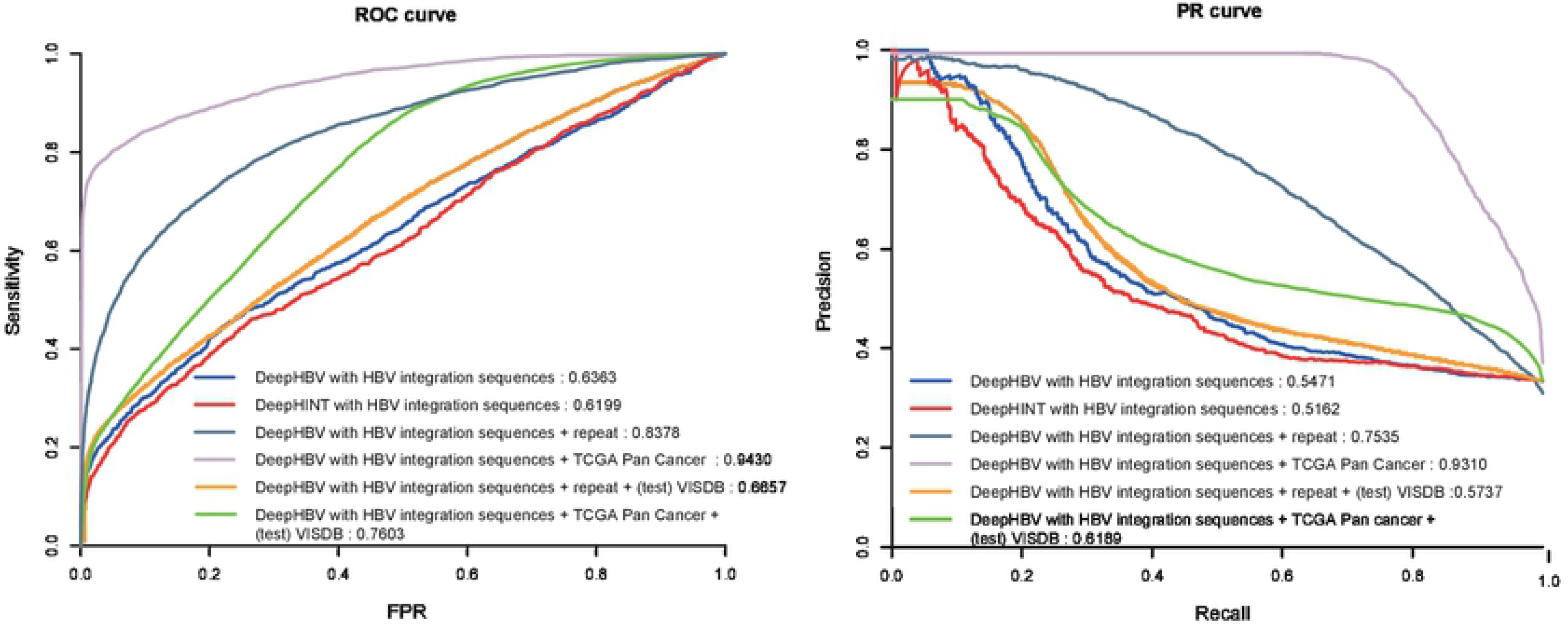
Attention intensive regions highlighted essential local genomic features on predicting HBV integration sites. Representative examples showed the positional relationship between the attention intensive sites and several genomic features using DeepHBV with HBV integration sequences + TCGA Pan Cancer model on (a) chr5:1,294,063-1,296,063 (hg38), (b) chr5: 1291277-1293277 (hg38). Each of these two sequences contains HBV integration sites from both dsVIS and VISDB. Enriched DNA binding proteins detected by HOMER from the attention intensive regions using the output of DeepHBV then we applied FIMO [1] to find the enriched motif position and label the motifs on attention intensive regions. UCSC genome browser [2] and Matplotlib [3] was used for visualisation. “HPV integration site” refers to the sites selected from our unpublished database used as testing samples. “Attention Intensive Sites” denotes the sites with top 5% attention weight. “RepeatMasker”, “TCGA Pan Cancer”, “DNase Clusters”, “Con20mammals”, “GeneHancer”, “Layered H3K27ac”, “Layered H3K36me3” are genomic features.

Convolution and pooling module will learn the patterns with translation invariance in deep learning, based on that deep learning network tend to learn the domains happened recurrently among different samples in the same pooling matrix, even if the learned feature was not at the same position in these different samples [20,21]. Attention intensive regions are more likely to be conserved due to the translation invariance in convolution and pooling module, and would give hints to the selection preference of HBV integration sites. Transcriptional factor-binding sites (TFBS) motifs are conserved genomic elements which can be critical to the regulation of downstream genes. Therefore, we tested whether TFBS played important roles in HBV integration preference. We used all HBV integration samples whose prediction scores were higher than 0.95 from dsVIS and VISDB separately to enrich local TFBS motifs in attention intensive regions by HOMER v 4.11.1 [22] with its vertebrates transcription factor databases (Table 1). From the result of DeepHBV with HBV integration sequences + TCGA Pan Cancer, binding sites of AR-halfsite, Arnt, Atf1, bHLHE40, bHLHE41, BMAL1, CLOCK, c-Myc, COUP-TFII, E2A, EBF1, Erra, Foxo3, HEB, HIC1, HIF-1b, LRF, Meis1, MITF, MNT, MyoG, n-Myc, NPAS2, NPAS, Nr5a2, Ptf1a, Snail1, Tbx5, Tbx6, TCF7, TEAD1, TEAD3, TEAD4, TEAD, Tgif1, Tgif2, THRb, USF1, Usf2, Zac1, ZEB1, ZFX, ZNF692, ZNF711 can be both enriched in attention intensive regions of dsVIS and VISDB sequences. We selected two representative samples to obtain a more intuitive display. Genomic features, HBV integration sites from dsVIS and VISDB, attention intensive sites and TFBS were aligned and shown in hg38 reference genome (Fig 4). Most attention intensive sites can be mapped to enrich TF motifs. And the clusters of high attention weight from the output of DeepHBV with HBV integration sites plus TCGA Pan Cancer showed the binding site of a tumour suppressor gene HIC1, circadian clock related elements BMAL1, CLOCK, c-Myc and NAPS2 (Fig 4). The data provided novel insights into HBV integration site selection preference and reveal biological importance that warrants future experimental confirmation.

**Table 1.**
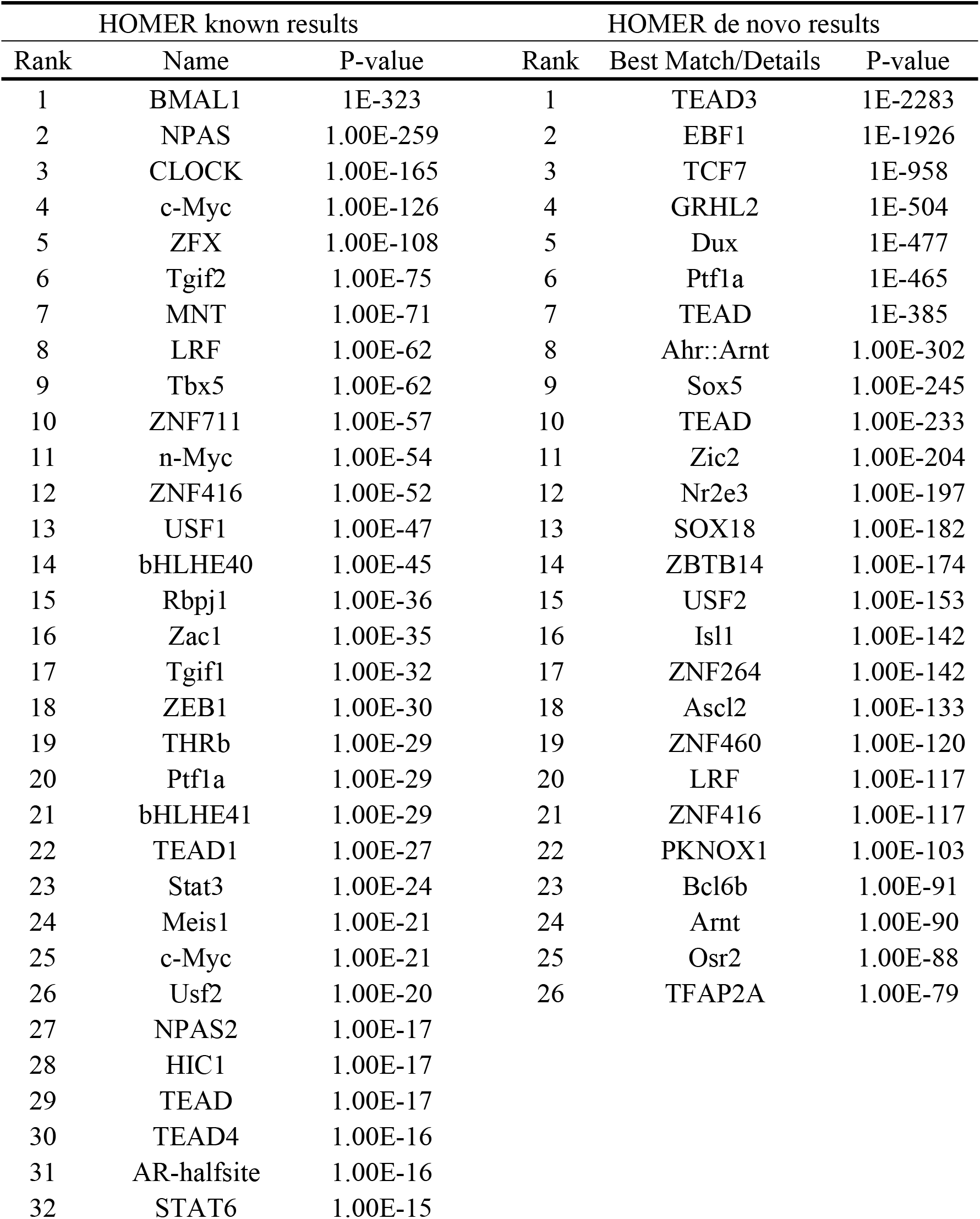
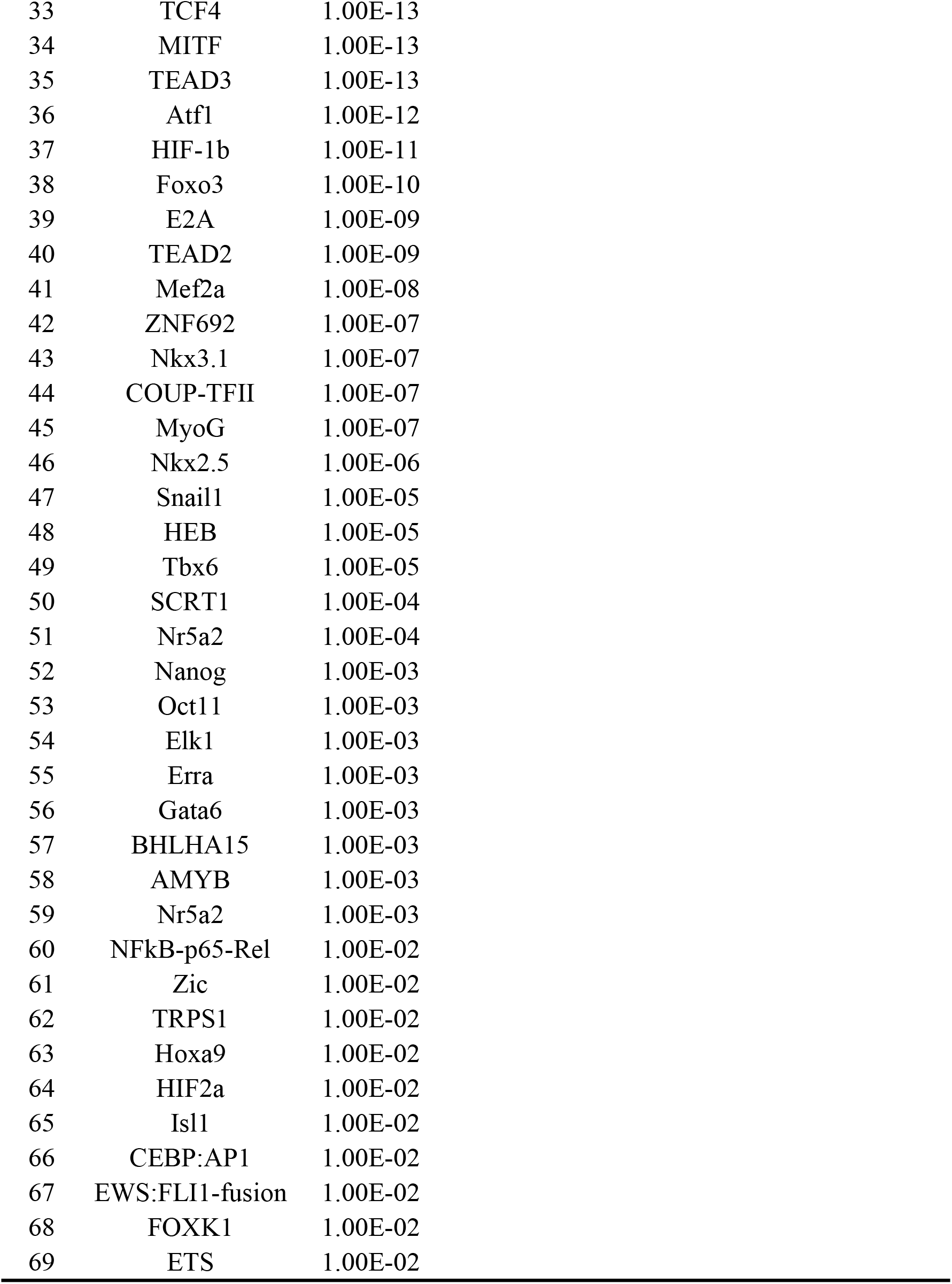
Enriched TFBS from attention intensive regions of DeepHBV with HBV integration sites + TCGA Pan Cancer peaks.

| HOMER known results |  |  | HOMER de novo results |  |  |
| --- | --- | --- | --- | --- | --- |
| Rank | Name | P-value | Rank | Best Match/Details | P-value |
| 1 | BMAL1 | 1E-323 | 1 | TEAD3 | 1E-2283 |
| 2 | NPAS | 1.00E-259 | 2 | EBF1 | 1E-1926 |
| 3 | CLOCK | 1.00E-165 | 3 | TCF7 | 1E-958 |
| 4 | c-Myc | 1.00E-126 | 4 | GRHL2 | 1E-504 |
| 5 | ZFX | 1.00E-108 | 5 | Dux | 1E-477 |
| 6 | Tgif2 | 1.00E-75 | 6 | Ptf1a | 1E-465 |
| 7 | MNT | 1.00E-71 | 7 | TEAD | 1E-385 |
| 8 | LRF | 1.00E-62 | 8 | Ahr::Arnt | 1.00E-302 |
| 9 | Tbx5 | 1.00E-62 | 9 | Sox5 | 1.00E-245 |
| 10 | ZNF711 | 1.00E-57 | 10 | TEAD | 1.00E-233 |
| 11 | n-Myc | 1.00E-54 | 11 | Zic2 | 1.00E-204 |
| 12 | ZNF416 | 1.00E-52 | 12 | Nr2e3 | 1.00E-197 |
| 13 | USF1 | 1.00E-47 | 13 | SOX18 | 1.00E-182 |
| 14 | bHLHE40 | 1.00E-45 | 14 | ZBTB14 | 1.00E-174 |
| 15 | Rbpj1 | 1.00E-36 | 15 | USF2 | 1.00E-153 |
| 16 | Zac1 | 1.00E-35 | 16 | Isl1 | 1.00E-142 |
| 17 | Tgif1 | 1.00E-32 | 17 | ZNF264 | 1.00E-142 |
| 18 | ZEB1 | 1.00E-30 | 18 | Ascl2 | 1.00E-133 |
| 19 | THRb | 1.00E-29 | 19 | ZNF460 | 1.00E-120 |
| 20 | Ptf1a | 1.00E-29 | 20 | LRF | 1.00E-117 |
| 21 | bHLHE41 | 1.00E-29 | 21 | ZNF416 | 1.00E-117 |
| 22 | TEAD1 | 1.00E-27 | 22 | PKNOX1 | 1.00E-103 |
| 23 | Stat3 | 1.00E-24 | 23 | Bcl6b | 1.00E-91 |
| 24 | Meis1 | 1.00E-21 | 24 | Arnt | 1.00E-90 |
| 25 | c-Myc | 1.00E-21 | 25 | Osr2 | 1.00E-88 |
| 26 | Usf2 | 1.00E-20 | 26 | TFAP2A | 1.00E-79 |
| 27 | NPAS2 | 1.00E-17 |  |  |  |
| 28 | HIC1 | 1.00E-17 |  |  |  |
| 29 | TEAD | 1.00E-17 |  |  |  |
| 30 | TEAD4 | 1.00E-16 |  |  |  |
| 31 | AR-halfsite | 1.00E-16 |  |  |  |
| 32 | STAT6 | 1.00E-15 |  |  |  |

|  |  |  |
| --- | --- | --- |
| 33 | TCF4 | 1.00E-13 |
| 34 | MITF | 1.00E-13 |
| 35 | TEAD3 | 1.00E-13 |
| 36 | Atf1 | 1.00E-12 |
| 37 | HIF-1b | 1.00E-11 |
| 38 | Foxo3 | 1.00E-10 |
| 39 | E2A | 1.00E-09 |
| 40 | TEAD2 | 1.00E-09 |
| 41 | Mef2a | 1.00E-08 |
| 42 | ZNF692 | 1.00E-07 |
| 43 | Nkx3.1 | 1.00E-07 |
| 44 | COUP-TFII | 1.00E-07 |
| 45 | MyoG | 1.00E-07 |
| 46 | Nkx2.5 | 1.00E-06 |
| 47 | Snail1 | 1.00E-05 |
| 48 | HEB | 1.00E-05 |
| 49 | Tbx6 | 1.00E-05 |
| 50 | SCRT1 | 1.00E-04 |
| 51 | Nr5a2 | 1.00E-04 |
| 52 | Nanog | 1.00E-03 |
| 53 | Oct11 | 1.00E-03 |
| 54 | Elk1 | 1.00E-03 |
| 55 | Erra | 1.00E-03 |
| 56 | Gata6 | 1.00E-03 |
| 57 | BHLHA15 | 1.00E-03 |
| 58 | AMYB | 1.00E-03 |
| 59 | Nr5a2 | 1.00E-03 |
| 60 | NFkB-p65-Rel | 1.00E-02 |
| 61 | Zic | 1.00E-02 |
| 62 | TRPS1 | 1.00E-02 |
| 63 | Hoxa9 | 1.00E-02 |
| 64 | HIF2a | 1.00E-02 |
| 65 | Isl1 | 1.00E-02 |
| 66 | CEBP:AP1 | 1.00E-02 |
| 67 | EWS:FLI1-fusion | 1.00E-02 |
| 68 | FO XK1 | 1.00E-02 |
| 69 | ETS | 1.00E-02 |

## Discussion

In this study, we developed an explainable attention-based deep learning model DeepHBV to predict HBV integration sites. In the comparison of DeepHBV and DeepHINT on predicting HBV integration sites (S3 Table), DeepHBV out-performed DeepHINT after adding genomic features due to its more suitable model structure and parameters on recognising the surroundings of HBV integration sites. We applied two convolution layers (1^st^ layer: 128 convolution kernels and the kernel size is 8; 2^nd^ layer: 256 convolution kernels and the kernel size is 6) and one pooling layer (with pooling size of 3) in DeepHBV while in DeepHINT the model only have one convolution layer (64 convolution kernels and the kernel size is 6) and one pooling layer (with pool size of 3). The increasing of convolution layers enables the information from higher dimensions can be extracted, the increasing of convolution kernels enables more feature information to be extracted [23].

We trained the DeepHBV model using three strategies (1) DNA sequences near HBV integration sites (HBV integration sequences), (2) HBV integration sequences + TCGA Pan Cancer peaks, (3) HBV integration sequences + repeat peaks. We found that the model with HBV integration sequences adding TCGA Pan Cancer or repeat can both significantly improve the model performance. And the DeepHBV with HBV integration sequences adding TCGA Pan Cancer peaks performed better on independent test dataset VISDB. However, the attention intensive regions cannot be well aligned to these genomic features. Thus, we further inferred that other features such as TFBS motifs may influence DeepHBV in the prediction process. And HOMER was applied to recognise these TFBS that might be related to HBV-related diseases or cancer development.

We noticed that the attention intensive regions identified by attention mechanism of DeepHBV with HBV integration sequences + TCGA Pan Cancer showed strong concentration on the binding site of the tumour suppressor gene HIC1, circadian clock-related elements BMAL1, CLOCK, c-Myc, NAPS2, and the transcription factors TEAD and Nr5a2. These DNA binding proteins were closely related to tumour development [24–30]. For instance, HIC1 is a tumour suppressor gene in hepatocarcinogenesis development [24,25]. BMAL1, CLOCK, c-Myc, NAPS2 all participate in the regulation of circadian clock [26], which is reported to promote HBV-related diseases [27,28]. In accordance, the binding motif of circadian clock-related elements were also enriched from the attention intensive regions of DeepHBV with HBV integration sequences + repeats, further confirming the results (S4 Table). In addition, the other transcription factors identified by Deep HBV are TEAD and Nr5a2. TEAD deregulation affected well-established cancer genes such as BRAF, KRAS, MYC, NF2 and LKB1, and showed high correlation with clinicopathological parameters in human malignancies [29]. Nr5a2 (also known as Liver receptor homolog-1, LRH-1) binds to the enhancer II (ENII) of HBV genes, and serves as a critical regulator of their expression [30].

In summary, DeepHBV is a robust deep learning model of using convolutional neural network to predict HBV integrations. Our data provide new insight into the preference for HBV integration and mechanism research on HBV induced cancer.

## Methods

### Data preparation

A detailed step-by-step instruction of DeepHBV was provided in S1 and S2 Notes. To obtain positive training and testing samples for DeepHBV, we extracted 1000 bp DNA sequences from upstream and 1000 bp DNA sequences from downstream of HBV integration sites as positive dataset, each sample was denoted as *S* = (*n*_1_,*n*_2_,…,*n*_2000_), where *n*_i_ represents the nucleotide in position *i*. DeepHBV, as a deep learning network also require negative samples that do not contain HBV integration sites as background area. The existing of HBV integration hot spots which contains several integration events within 30~100 kb range [13] prompted us that we should selected background area keeping enough distance from known HBV integration sites. Thus, we discarded the regions around known HBV integration sites with length 50 kb on hg38 reference genome and selected 2 kb length DNA sequences randomly on remained regions as negative samples.

We encoded extracted DNA sequences using one-hot code to make the calculation of distance between features in training and the calculation of similarity more accuracy. Original DNA sequences were converted to binary matrices of 4-bit length where each dimension corresponds to one nucleotide type. Finally, we converted a 2000 bp DNA sequence into a 2000×4 binary matrix.

### Feature extraction

DeepHBV model first applied convolution and pooling module to learn and obtain sequence features around HBV integration sites (S1 Fig). Each binary matrix representing a DNA sequence entered the convolution and pooling module to execute convolution calculation. We employed multiple variant convolution kernels to calculation in order to obtain different features. S = (*n*_1_,*n*_2_,…,*n*_2000_) denoted as a specific DNA sequence and E represented the binary matrix-encoded from S, the convolutional calculation in convolution layer refers to *X* = *conv*(*E*), which can be described as:

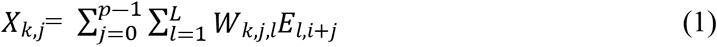

Where 1 ≤ *k* ≤ *d*, *d* refers to the number of kernels, 1 ≤ *i* ≤ *n* ― *p* +1, *i* refers to the index, *p* refers to the kernel size, n refers to input sequence length, *W* refers to the kernel weight.

Convolutional layer activated eigen vectors using Rectified Linear Unit (ReLU) after extracting relative eigen vectors. ReLU is an activation function in artificial neural networks which can be described as *f*(*x*) = max (0,*x*). We applied ReLU on the output matrix of each convolution layer and mapped each element on a sparse matrix. ReLU imitates real neuron activation, which enables data fitted to the model better. Then we applied max-pooling strategy to complete dimension reduction as well as support the maximum retention of predicted information. Till now, we achieved the final eigen vector *F_c_* from the binary matrix represented DNA sequence after feature extracting in convolution and pooling module.

### Attention mechanism in DeepHBV model

DeepHBV added attention mechanism in order to capture and understand the position contribution in abstracted eigen-vector *F_c_*. Eigen-vector entered the attention layer, which will calculate a weight value to each dimension in *F_c_*. The attention weight represents the contribution level of the convolutional neural network (CNN) in that position. The output of attention weight *t_j_* is the contribution score, larger *t_j_* score means bigger contribution in this position to HBV integration sites prediction. All contribution scores were normalized to achieve the dense eigenvector matrix, which denoted as *F_a_*:

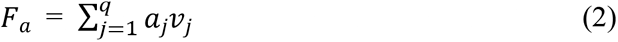

Where,

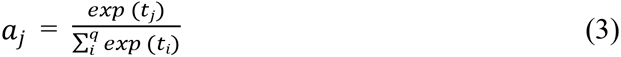

Where *a_j_* represents the relevant normalisation score, *v_j_* represents the eigenvector at position *j* of the input eigenmatrix. Each position represents an extracted eigen-vector in each convolution kernel.

The convolution-pooling module and the attention mechanism module need to be combined in model prediction progress, in another word, eigen-vector *F_c_* and relative eigen important score *F_a_* should work together in HBV integration sites prediction.

We linked the values in eigen-vector *F_c_* and linearly mapped them to a new vector *F_v_*, which is:

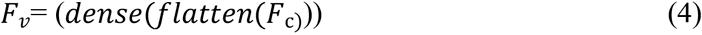

In this step, flatten layer performed function *flatten*() to reduce dimension and concatenate data; function *dense*() was executed by dense layer, which will map dimension-reduced data to a single value. Then *F_v_* and *F_a_* concatenated vector entered linear classifier prediction to calculate the probability of HBV integration happened within the current sequence, with:

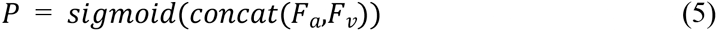

Where *P* is the predicted score, *sigmoid*() represents the activation function acted as classifier in final output, *concat* () represents the concatenate operation.

In the meantime, if we give the output eigenvector *F_c_* from convolution-and-pooling module as input, and execute attention mechanism, weight vector *W* can be achieved:

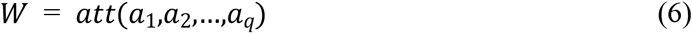

Where *att*() refers to the attention mechanism, *a_i_* denotes the eigenvector in *i^th^* dimension in the eigenmatrix *W*, represents the dataset containing contribution scores of each position in the eigenmatrix extracted by convolution-and-pooling module.

### DeepHBV model training

After confirming each parameter in DeepHBV (S1 Table), we trained the deep learning neural network model DeepHBV via binary crossentropy. The loss function of DeepHBV can be defined as:

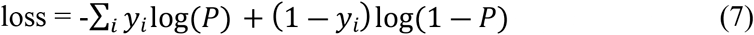

Where, *y_i_* is the prediction score *P*, is the binary tag value of that sequence (in this dataset, positive samples were labelled as 1 and negative samples were labelled as 0). Back propagation algorithm was adapted in training progress and Nesterov-accelerated adaptive moment estimation (Nadam) gradient descent algorithm was applied to optimise parameter initialization.

The deep learning neural network model adapted Python 3.7, Keras library 2.2.4 [14] using three NVIDIA® Tesla V100-PCIE-32G (NVIDIA Corporation, California, USA) for training and testing. DeepHBV takes around 90 min and 30 s for model training and testing respectively using the computational platform under such software and hardware settings.

## Data Availability

DeepHBV is available as an open-source software and can be downloaded from https://github.com/JiuxingLiang/DeepHBV.git

## Supporting information

**S1 Fig. DeepHBV framework.** Each part represents a layer in neural network and *n* × *n* stands for the output dimension which was explained in S2 Note. Two continuous convolution layers were used to extract features; max-pooling layers can reduce the dimension while keeping the feature matrix has the ability to predicting information; dropout layer randomly drop some results to prevent over-fit; flatten layer is responsible for reduce the dimensions and connect them; dense layer is used to map the output from last layer to a specific value; attention layer and attention flatten are used to give a weight score to each dimension in the feature matrix; concatenate layer concatenates captured features and importance scores of those features from the convolution module and the attention mechanism model. Prediction Output offered the final output reveals the probability of HBV infection.

**S2 Fig. Prediction performance on the HBV integration dataset with different types of genomic features added in.** We found that character 1 and character 3 outperformed the DeepHBV model with an significant increase in AUPR and AUROC score on character 1 and character 3, indicating that DeepHBV can capture genomic features from character 1 and character 3 effectively, so we did further analysis on each single items in character group 1 and 3, and found that Repeats and TCGA Pan Cancer are the genomic features that can be captured by DeepHBV which significantly improved model performance. DeepHBV with HBV integration sequences + repeats reached the AUROC of 0.8378 and the AUPR of 0.7535, which DeepHBV with HBV integration sequences + TCGA Pan Cancer reached the AUROC of 0.9430 and the AUPR of 0.9310.

**S1 Table. The parameters for the deep neural network used in DeepHBV.**

**S2 Table. Genomic features and sources. (Access date: Novemember 16^th^, 2019)**

**S3 Table. Comparison of DeepHBV and DeepHINT result record.**

**S4 Table. Enriched TFBS from attention intensive regions of DeepHBV with HBV integration sites + repeat peaks.**

**S1 Note. DeepHBV framework.** DeepHBV neural network structure design and hyperparameters involved in DeepHBV are noted.

**S2 Note. Mathematical matters of the DeepHBV.** There are explanations for 8 mathematical matters (i.e. encoding DNA sequences, convolution layers, the max pooling layer, dropout layer, attention layer, concatenate layer, linear classifier and optimisation algorithm) of the DeepHBV in this part.

## Reference

1. Liang TJ. Hepatitis B: the virus and disease. Hepatology 2009;49(5 Suppl):S13–21.

2. Tu T, Budzinska MA, Shackel NA et al. HBV DNA Integration: Molecular Mechanisms and Clinical Implications. Viruses 2017;9(4).

3. Sung WK, Zheng H, Li S et al. Genome-wide survey of recurrent HBV integration in hepatocellular carcinoma. Nat Genet 2012;44(7):765–9.

4. Zhao LH, Liu X, Yan HX et al. Genomic and oncogenic preference of HBV integration in hepatocellular carcinoma. Nat Commun 2016;7:12992.

5. Ding D, Lou X, Hua D et al. Recurrent targeted genes of hepatitis B virus in the liver cancer genomes identified by a next-generation sequencing-based approach. PLoS Genet 2012;8(12):e1003065.

6. Tu T, Budzinska MA, Vondran FWR et al. Hepatitis B Virus DNA Integration Occurs Early in the Viral Life Cycle in an In Vitro Infection Model via Sodium Taurocholate Cotransporting Polypeptide-Dependent Uptake of Enveloped Virus Particles. J Virol 2018;92(11).

7. Mason WS, Gill US, Litwin S et al. HBV DNA Integration and Clonal Hepatocyte Expansion in Chronic Hepatitis B Patients Considered Immune Tolerant. Gastroenterology 2016;151(5):986–998 e4.

8. Litjens G, Kooi T, Bejnordi BE et al. A survey on deep learning in medical image analysis. Med Image Anal 2017;42:60–88.

9. Bailey TL, Baker ME, Elkan CP. An artificial intelligence approach to motif discovery in protein sequences: Application to steroid dehydrogenases. The Journal of Steroid Biochemistry and Molecular Biology 1997;62(1):29–44.

10. Yamashita R, Nishio M, Do RKG et al. Convolutional neural networks: an overview and application in radiology. Insights into Imaging 2018;9(4):611–629.

11. Bahdanau D, Cho K, Bengio Y. Neural Machine Translation by Jointly Learning to Align and Translate. Computer Science 2014.

12. Guidotti R, Monreale A, Ruggieri S et al. A Survey of Methods for Explaining Black Box Models. ACM Comput. Surv. 2018;51(5):Article 93.

13. Hu Z, Zhu D, Wang W et al. Genome-wide profiling of HPV integration in cervical cancer identifies clustered genomic hot spots and a potential microhomology-mediated integration mechanism. Nat Genet 2015;47(2):158–63.

14. Chollet Fao. Keras. 2015.

15. Hu H, Xiao A, Zhang S et al. DeepHINT: understanding HIV-1 integration via deep learning with attention. Bioinformatics 2019;35(10):1660–1667.

16. Haeussler M, Zweig AS, Tyner C et al. The UCSC Genome Browser database: 2019 update. Nucleic Acids Res 2019;47(D1):D853–D858.

17. Inoue F, Kircher M, Martin B et al. A systematic comparison reveals substantial differences in chromosomal versus episomal encoding of enhancer activity. Genome Res 2017;27(1):38–52.

18. Robinson JT, Thorvaldsdottir H, Winckler W et al. Integrative genomics viewer. Nature Biotechnology 2011;29(1):24–26.

19. Tang D, Li B, Xu T et al. VISDB: a manually curated database of viral integration sites in the human genome. Nucleic Acids Res 2019.

20. Zhang W, Itoh K, Tanida J et al. Parallel distributed processing model with local space-invariant interconnections and its optical architecture. Appl Opt 1990;29(32):4790–7.

21. Bruna J, Zaremba W, Szlam A et al. Spectral Networks and Locally Connected Networks on Graphs. Computer Science 2013.

22. Heinz S, Benner C, Spann N et al. Simple Combinations of Lineage-Determining Transcription Factors Prime cis-Regulatory Elements Required for Macrophage and B Cell Identities. Molecular Cell 2010;38(4):576–589.

23. Seide F, Gang L, Dong Y. Conversational speech transcription using context-dependent deep neural networks. 2012.

24. Taniguchi K, Roberts LR, Aderca IN et al. Mutational spectrum of beta-catenin, AXIN1, and AXIN2 in hepatocellular carcinomas and hepatoblastomas. Oncogene 2002;21(31):4863–71.

25. Zheng J, Xiong D, Sun X et al. Signification of Hypermethylated in Cancer 1 (HIC1) as Tumor Suppressor Gene in Tumor Progression. Cancer Microenviron 2012;5(3):285–93.

26. Paibomesai MI, Moghadam HK, Ferguson MM et al. Clock genes and their genomic distributions in three species of salmonid fishes: Associations with genes regulating sexual maturation and cell cycling. BMC Res Notes 2010;3:215.

27. Fekry B, Ribas-Latre A, Baumgartner C et al. Incompatibility of the circadian protein BMAL1 and HNF4alpha in hepatocellular carcinoma. Nat Commun 2018;9(1):4349.

28. Mukherji A, Bailey SM, Staels B et al. The circadian clock and liver function in health and disease. J Hepatol 2019;71(1):200–211.

29. Huh HD, Kim DH, Jeong HS et al. Regulation of TEAD Transcription Factors in Cancer Biology. Cells 2019;8(6).

30. Cai YN, Zhou Q, Kong YY et al. LRH-1/hB1F and HNF1 synergistically up-regulate hepatitis B virus gene transcription and DNA replication. Cell Research 2003;13(6):451–458.

## References

1. Grant CE, Bailey TL, Noble WS. FIMO: scanning for occurrences of a given motif. Bioinformatics 2011;27(7):1017–8.

2. Haeussler M, Zweig AS, Tyner C et al. The UCSC Genome Browser database: 2019 update. Nucleic Acids Res 2019;47(D1):D853–D858.

3. Hunter JD. Matplotlib: A 2D Graphics Environment. Computing in Science & Engineering 2007;9(3):90–95.

